# Genome of a Giant (Trevally): *Caranx ignobilis*

**DOI:** 10.1101/2021.09.11.459923

**Authors:** Brandon D. Pickett, Jessica R. Glass, Perry G. Ridge, John S. K. Kauwe

**Affiliations:** Department of Biology, Brigham Young University, Provo, Utah, USA; South African Institute for Aquatic Biodiversity, Makhanda, South Africa; College of Fisheries and Ocean Sciences, University of Alaska Fairbanks, Fairbanks, Alaska, USA; Brigham Young University – Hawai‘i, Laie, Hawai‘i, USA

**Keywords:** Giant Trevally, Kingfish, ‘Ulua, Carangidae, Carangiformes, *de novo* genome assembly

## Abstract

*Caranx ignobilis*, commonly known as the kingfish or giant trevally, is a large, reef-associated apex predator. It is a prized sportfish, targeted heavily throughout its tropical and subtropical range in the Indian and Pacific Oceans, and it has drawn significant interest in aquaculture due to an unusual tolerance for freshwater. In this study, we present a high-quality nuclear genome assembly of a *C. ignobilis* individual from Hawaiian waters, which have recently been shown to host a genetically distinct population. The assembly has a contig NG50 of 7.3Mbp and scaffold NG50 of 46.3Mbp. Twenty-five of the 203 scaffolds contain 90% of the genome. We also present the raw Pacific Biosciences continuous long-reads from which the assembly was created. A Hi-C dataset (Dovetail Genomics Omni-C) and Illumina-based RNA-seq from eight tissues are also presented; the latter of which can be particularly useful for annotation and studies of freshwater tolerance. Overall, this genome assembly and supporting data is a valuable tool for ecological and comparative genomics studies of kingfish and other carangoid fishes.

## BACKGROUND & SUMMARY

The “genomic revolution” continues to rapidly advance our understanding of human evolution, as well as the evolution of non-model organisms ^1^. Comparative genomic approaches using whole-genome datasets allow for discoveries at every scale: from genome to chromosome to organism to entire clades of organisms. Genomic datasets for non-model marine teleost fishes, the most diverse clade of vertebrates, are invaluable for investigating evolutionary questions relating to adaptation, selection, genome duplication, and phylogenetic conservatism in vertebrates.

We present a high-quality genome assembly of the marine teleost, giant trevally (*Caranx ignobilis;* Carangiformes: Carangoidei; Fig. 1). This assembly serves as a valuable resource for the field of evolutionary biology, ecology, and phylogenetics. *Caranx ignobilis* is a member of the Carangini clade, the most specious subclade within Carangoidei. Carangoid fishes are known for their extreme diversity in morphology and ecology ^2,3^. The giant trevally, specifically, is known to be highly tolerant of freshwater environments, leading to a surge of interest in this species for aquaculture ^4–6^ and making it an ideal candidate species to investigate linkages between genotype and phenotype in the context of freshwater adaptation by marine fishes ^7,8^. *Caranx ignobilis* is an apex predator in tropical and subtropical reefs and coastal environments in the Indian and Pacific Oceans ^9^ and is heavily targeted by small-scale and recreational fisheries throughout its range. Understanding its evolutionary and ecological role in ecosystem structure and function is important for fisheries management and the protection of reef and coral ecosystems. Importantly, new putative populations of *C. ignobilis* in the Indian and Pacific Oceans have recently been described using genomic datasets ^10^. A high-quality genome thus allows for the inference of demographic history, genomic signals of selection and adaption, and comparative genomic studies with other Carangoid fishes, such as hybridization with the closely related bluefin trevally, *Caranx melampygus* ^11^.

**Figure 1.**
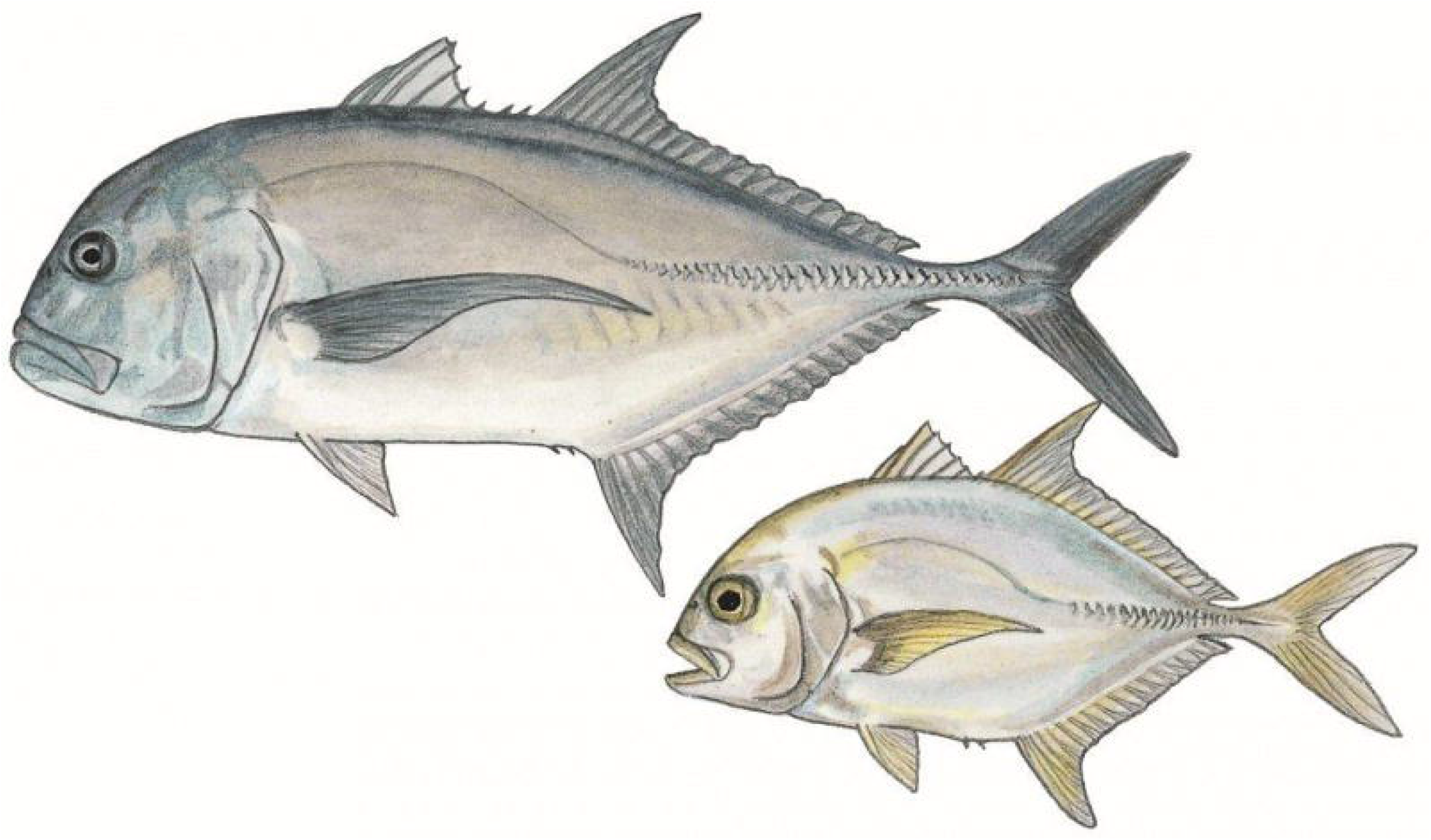
Giant trevally (*Caranx ignobilis*) adult and juvenile. Illustration by Elaine Heemstra, courtesy of the South African Institute for Aquatic Biodiversity.

For this *C. ignobilis* assembly, we present results using 58.25 Gbp of Pacific Biosciences (PacBio) Single-molecule, Real-time (SMRT) sequencing data. Illumina paired-end sequencing data was also generated with libraries for both RNA-seq and Hi-C, totaling 347.6 Gbp. Both were used for scaffolding purposes and are valuable datasets individually. The estimated genome size was 625.92 Mbp ^12,13^, of which 96.7% is covered by known bases in the primary haploid assembly. In addition to being highly contiguous, the genome assembly contained complete, unduplicated copies of >95% of expected single-copy orthologs, suggesting the assembly is reasonably complete. The assembly and supporting sequencing datasets are sufficiently high-quality to serve as a valuable resource for a variety of prospective comparative and population genomics studies.

## METHODS

An overview of the methods used in this study is provided here. Where appropriate, additional details, such as the code for custom scripts and the commands used to run software, are provided in the Supplementary Bioinformatics Methods (Supplementary File 1).

### Sample Acquisition & Sequencing

Blood, brain, eye, fin, gill, heart, kidney, liver, and muscle tissues from one *C. ignobilis* individual were collected off the coast of O’ahu (near Kaneohe, Hawai’i, USA) in April 2019. Blood was preserved in EDTA, and other tissue samples were flash-frozen in liquid nitrogen. All samples were packaged in dry ice for transportation to Brigham Young University (BYU; Provo, Utah, USA) for storage at −80°C until sequencing. The blood sample was used to create the Omni-C dataset. All non-blood tissue samples were used for short-read RNA sequencing; the heart tissue was also used for long-read DNA sequencing.

DNA was prepared for long-read sequencing with a Pacific Biosciences (PacBio; Menlo Park, California, USA; https://www.pacb.com) SMRTbell Library kit, adhering to the following protocol: “Procedure & Checklist - Preparing gDNA Libraries Using the SMRTbell Express Template Preparation Kit 2.0”. Continuous long-read (CLR) sequencing was performed on seven SMRT cells for a 10-hour movie on the PacBio Sequel at the BYU DNA Sequencing Center (DNASC; https://dnasc.byu.edu), a PacBio Certified Service Provider. RNA was prepared with Roche (Basel, Switzerland; https://sequencing.roche.com) KAPA Stranded RNA-seq kit, following recommended protocols. Paired-end sequencing was performed in High Output mode for 125 cycles with the eight samples across two lanes on the Illumina (San Diego, California, USA; https://www.illumina.com) Hi-Seq 2500 at the DNASC. Finally, the “Omni-C Proximity Ligation Assay Protocol” version 1.0 was followed using a Dovetail Genomics Omni-C kit to prepare for Illumina Paired-end sequencing. Adapters were provided by Integrated DNA Technologies, and sequencing proceeded in Rapid Run mode for 250 cycles in one lane on an Illumina Hi-Seq 2500.

### Sequence Assembly, Duplicate Purging, and Scaffolding

The PacBio CLR reads were self-corrected and assembled with Canu v1.8 ^14^. To get a haploid representation of the genome, duplicates were purged with purge_dups v1.2.5 ^15^. The primary set of 329 contigs was selected for scaffolding with Omni-C data, which required reads to be mapped to the assembly before determining how to order and orient the contigs. The Omni-C reads were aligned following the Arima Genomics (San Diego, California, USA; https://arimagenomics.com) Mapping Pipeline commit #2e74ea4 (https://github.com/ArimaGenomics/mapping_pipeline), which relied on BWA-MEM2 v2.1 ^16,17^, Picard v2.19.2 ^18^, and SAMtools v1.9 ^19^. BEDTools v2.28.0 ^20^ was used to prepare the Omni-C alignments for scaffolding with SALSA commit #974589f ^21^. Before the scaffolding step was performed, SALSA cleaned the assembly by breaking misassemblies as determined by Omni-C read mappings. This set of contigs was then used simultaneously for both the remainder of the SALSA pipeline and for scaffolding with Rascaf v1.0.2 commit #690f618 ^22^ using the RNA-seq data from all tissues aligned using HiSat v0.1.6-beta ^23^. The two sets of scaffolds were combined using custom Python (https://www.python.org) scripts, which used the Omni-C scaffolds as a starting point and added compatible joins from the RNA-seq evidence. Contamination was removed from the final set of scaffolds as identified during the NCBI submission process; all gaps were also adjusted to a fixed size (100 Ns). Repeat characterization was performed with RepeatMasker v4.1.2-p1 ^24^ using Dfam v3.3 ^25^ and the RepBase RepeatMasker Library v20181026 ^26,27^.

### Genome Assembly Validation

At each phase of the assembly, continuity statistics, e.g., N50 and auNG, were calculated with caln50 commit #3e1b2be (https://github.com/lh3/calN50) and a custom Python script (Table 3). The genome size (625.92 Mbp) provided to Canu and used for the computation of assembly statistics was based on the C-value of 0.64 from Hardie and Hebert ^12^ as recorded in the Animal Genome Size Database ^13^. Assembly completeness was also assessed at each phase using single-copy orthologs from the Actinopterygii set of OrthoDB v10 ^28^ as identified by BUSCO v4.0.6 ^29^ (Table 4). The scaffolds were visually inspected using a Hi-C contact matrix created with PretextView v0.1.4 (https://github.com/wtsi-hpag/PretextView) and PretextMap v0.1.4 (https://github.com/wtsi-hpag/PretextMap) with SAMtools v1.10 ^19^.

Visual comparisons with other carangoid genomes were created for cursory validation and observation of general synteny. Dot plots were generated using Mashmap v2.0 commit #ffeef48 ^30^ (-f ‘one-to-one’ --pi 95 -s 10000) and a comparison of single-copy orthologs was created using ChrOrthLink commit #d29b10b after assessment with BUSCO v3.0.6 ^29^ using the Vertebrata set from OrthoDB v9 ^31^. The genome assemblies obtained from NCBI for these analyses were the following (alphabetical order): *Caranx melampygus* (bluefin trevally) ^11^, *Echeneis naucrates* (live suckershark) ^32,33^, *Seriola dumerili* (greater amberjack) ^32,33^, *Seriola quinqueradiata* (yellowtail) ^34,35^, *Seriola rivoliana* (longfin yellowtail) ^36^, *Trachinotus ovatus* (golden pompano) ^37,38^, and *Trachurus trachurus* (Atlantic horse mackerel) ^39–41^.

## TECHNICAL VALIDATION

### Sequencing

Continuous long-read sequencing (PacBio) generated 3.74M reads with a total of 58.25 Gbp, which is approximately 93x physical coverage of the genome. The mean and N50 read lengths were 15,591.278 and 27,441, respectively. The longest read was 129,643bp. The read length distribution is plotted in Figure 2. A summary of the results for the sequencing run is available in Table 1. This genome represents the second for the *Caranx* genus and ranks highly in terms of N50 among available carangoid genomes ^38,40^.

**Figure 2.**
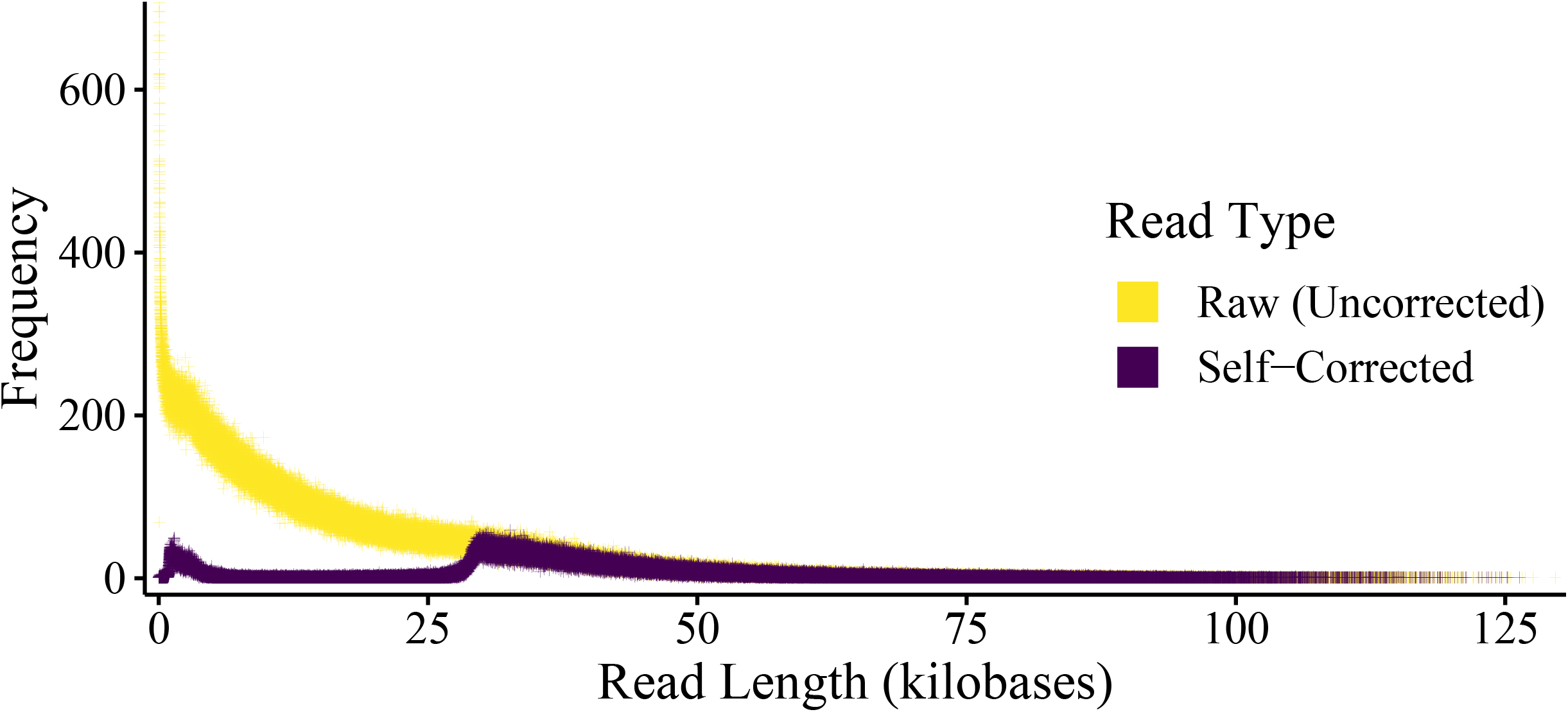
Frequency of Pacific Biosciences Read Lengths. The change in read length distribution is demonstrated as reads are corrected. The dramatic shift from raw to corrected reads is evident. Reads were corrected by consensus using the correction phase of Canu v1.8.

**Table 1.** Sequencing Information.

| Company | Illumina | Illumina | PacBio |
| --- | --- | --- | --- |
| **Instrument** | Hi-Seq 2500 | Hi-Seq 2500 | Sequel I |
| **Mode** | High Output | Rapid Run | NA |
| **Sequencing Type** | PE | Omni-C, PE | SMRT, CLR |
| **Duration** | 125 cycles | 250 cycles | 10 hours |
| **Specimen** | 1 | 1 | 1 |
| **Tissues** | Brain, Eye, Fin, Gill, Heart, Kidney, Liver, Muscle | Blood | Heart |
| **Molecule** | RNA | DNA | DNA |
| **Millions of Read( Pair)s** | 435.99 | 169.11 | 3.74 |
| **Mean Read Length<sub>(bp)</sub>** | 124.21 | 239.26 | 15,591.28 |
| **Read N50<sub>(bp)</sub>** | 125 | 250 | 27,441 |
| **Nucleotides<sub>(Gbp)</sub>** | 108.30 | 80.92 | 58.25 |

The results from each type of DNA and RNA sequencing from *Caranx ignobilis*. PE=Paired-end reads. SMRT=Single-Molecule, Real-Time sequencing. CLR=Continuous Long-reads.

RNA-seq from the eight tissues (i.e., brain, eye, fin, gill, heart, kidney, liver, and muscle) generated 435.99M pairs of reads totaling 108.30Gbp. Across all eight tissues, the mean and N50 read lengths were 124.21 and 125, respectively. The combined results from all eight tissues are represented in Table 1, while the results from each tissue are made available in Table 2. Omni-C sequencing generated 80.92 Gbp of data across 169.1M read pairs. The N50 and mean read length were respectively 250 and 239.3. The Omni-C results are also represented in Table 1 with the PacBio and RNA-seq data. The RNA-seq and Omni-C reads were not corrected, but the quality was assessed using fastqc ^42^.

**Table 2.** RNA Sequencing Details per Tissue.

|  | Millions<br>of Read<br>Pairs | Mean<br>Read<br>Length | Read<br>N50 | Nucleotides<br>(Gbp) |
| --- | --- | --- | --- | --- |
| Brain | 45.59 | 124.17 | 125 | 11.32 |
| Eye | 52.02 | 124.26 | 125 | 12.93 |
| Fin | 50.13 | 124.16 | 125 | 12.45 |
| Gill | 55.56 | 124.22 | 125 | 13.80 |
| Heart | 57.87 | 124.29 | 125 | 14.39 |
| Kidney | 58.73 | 124.16 | 125 | 14.58 |
| Liver | 58.25 | 124.23 | 125 | 14.47 |
| Muscle | 57.84 | 124.16 | 125 | 14.36 |
| **All** | **435.99** | **124.21** | **125** | **108.30** |

The results of RNA sequencing for each tissue from one *Caranx ignobilis* individual. The eight tissues were spread across two lanes and run on an Illumina Hi-Seq 2500 in Rapid Run mode for 250 cycles to generate paired-end reads. Unless otherwise specified, lengths of nucleotide sequences are measured in base pairs (bp).

### PacBio CLR Error Correction

The correction process reduced the number of reads from 3.74M to 656K and the total number of bases from 58.3Gbp to 23.9Gbp for an approximate physical coverage of 38.3x. The mean and N50 read lengths were changed from 15,591 and 27,441 to 36,475 and 40,065, respectively. The longest read was 126,321 bases. The distribution of corrected read lengths can be viewed relative to the raw read lengths in Figure 2.

### Genome Assembly, Duplicate Purging, and Scaffolding

The initial assembly from Canu was comprised of 1.8K contigs with a total assembly size of 758Mbp. This was a diploid assembly in the sense that both haplotypes were present and intermixed, separated whenever a bubble in the assembly graph prevented a single reasonable contig. Duplicate purging to get a haploid representation of the genome (albeit with inevitable haplotype switching) and fixing misassemblies with evidence from Hi-C data yielded 343 contigs with a total assembly size of 605Mbp. The mean contig length, N50, NG50, and maximum contig length were 1.8Mbp, 7.7Mbp, 7.3Mbp, and 19.6Mbp, respectively. The L50 was 23, and the LG50 was 25. The auNG was to 8.55M. These values represent modest reductions from the original Canu assembly (as expected), and they can be visualized in the area under the NG-curve as presented in Figure 3. (Also see Table 3)

**Figure 3.**
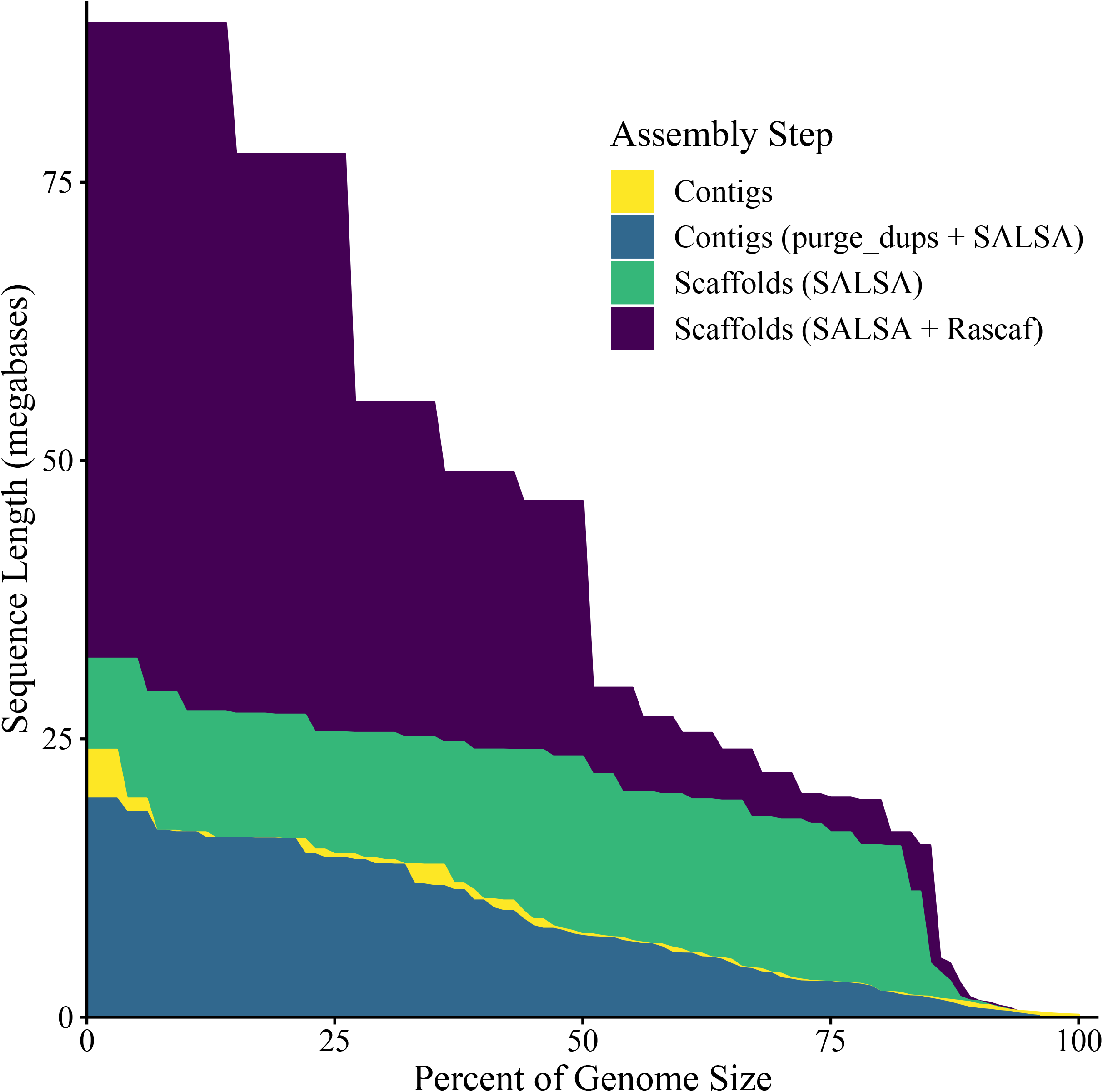
Area Under the NG-curve (auNG) for each Assembly Step. The NG-curve and the area under it are plotted for the contigs and scaffolds. This visually demonstrates an increase in continuity from contigs to scaffolds. Scaffolding with RNA-seq data – which has minimal effect on its own (data not shown) – further increases the scaffold-level continuity. This plot also shows that duplicate purging and fixing misassemblies slightly reduced contig-level continuity, which is expected.

**Table 3.** Continuity Statistics.

|  | Contigs | Contigs<br>(purge_dups) | Contigs<br>(purge_dups +<br>SALSA) | Scaffolds<br>(SALSA) | Scaffolds<br>(SALSA +<br>Rascaf) | Scaffolds |
| --- | --- | --- | --- | --- | --- | --- |
| **Sequences** | 1,804 | 329 | 343 | 240 | 209 | 203 |
| **Known**<br>**Bases** | 757.523 Mbp | 605.140 Mbp | 605.140 Mbp | 605.140 Mbp | 605.140 Mbp | 605.115 |
| **Mean**<br>**Length** | 0.420 Mbp | 1.839 Mbp | 1.764 Mbp | 2.521 Mbp | 2.895 Mbp | 2.981 Mbp |
| **Max.**<br>**Length** | 23.990 Mbp | 23.990 Mbp | 19.607 Mbp | 32.157 Mbp | 89.251 Mbp | 89.251 Mbp |
| **NG50** | 7.412 Mbp | 7.412 Mbp | 7.261 Mbp | 23.385 Mbp | 46.318 Mbp | 46.303 Mbp |
| **NG90** | 1.097 Mbp | 0.950 Mbp | 0.700 Mbp | 1.386 Mbp | 1.410 Mbp | 1.410 Mbp |
| **LG50** | 24 | 24 | 25 | 12 | 5 | 5 |
| **LG90** | 103 | 105 | 114 | 39 | 25 | 25 |
| **auNG** | 9.090 M | 9.051 M | 8.549 M | 19.716 M | 42.606 M | 42.600 M |
| **Sequences**<br>**with Gaps** | - | - | - | 40 | 35 | 35 |
| **Gaps** | - | - | - | 103 | 134 | 133 |
| **Unknown**<br>**Bases** | - | - | - | 51,500 | 52,027 | 13,300 |
| **Mean**<br>**Gap**<br>**Length** | - | - | - | 500 | 388.261 | 100 |

Continuity statistics for the *Caranx ignobilis* genome assembly at the contig and scaffold level. Note that the auNG value is the area under the NG curve, not the N curve. The final set of scaffolds (far right column) is the same as "Scaffolds (SALSA + Rascaf)" except that identified contaminants were manually removed from the assembly and gaps were unified to 100 Ns. Unless otherwise specified, all nucleotide sequences are measured in base pairs (bp).

Paired-end Illumina reads, such as those produced from Hi-C or RNA-seq libraries can provide information to order and orient contigs into scaffolds, but they contain insufficient information to utilize for gap-filling procedures. Accordingly, the result on assembly statistics should increase length, decrease the number of sequences, and leave the number of known bases unchanged. This pattern is evident in the assembly statistics from our iterative scaffolding procedure (Table 3). It is important to note that SALSA and Rascaf introduce gaps of unknown size, and they respectively use fixed runs of Ns of lengths 500 and 17 to represent such gaps. For submission to NCBI, these gaps were converted to a fixed length of 100 Ns, and the evidence for whether joins were supported by Hi-C data or RNA-seq data was submitted in an accompanying file in AGP format (https://www.ncbi.nlm.nih.gov/assembly/agp/AGP_Specification). The NCBI submission process also identified minor contaminants in some sequences, which were manually removed. The final set of scaffolds had an NG50 of 46.3Mbp and an auNG of 42.6M (Fig. 3; Table 3). All joins are represented in a contact matrix (Fig. 4), which shows the Hi-C evidence for the assembly. Some joins are poorly supported by the Hi-C evidence, which is not surprising as some joins were made by RNA-seq evidence instead. Without manual curation, it is difficult to ascertain whether any individual such join is spurious.

**Figure 4.**
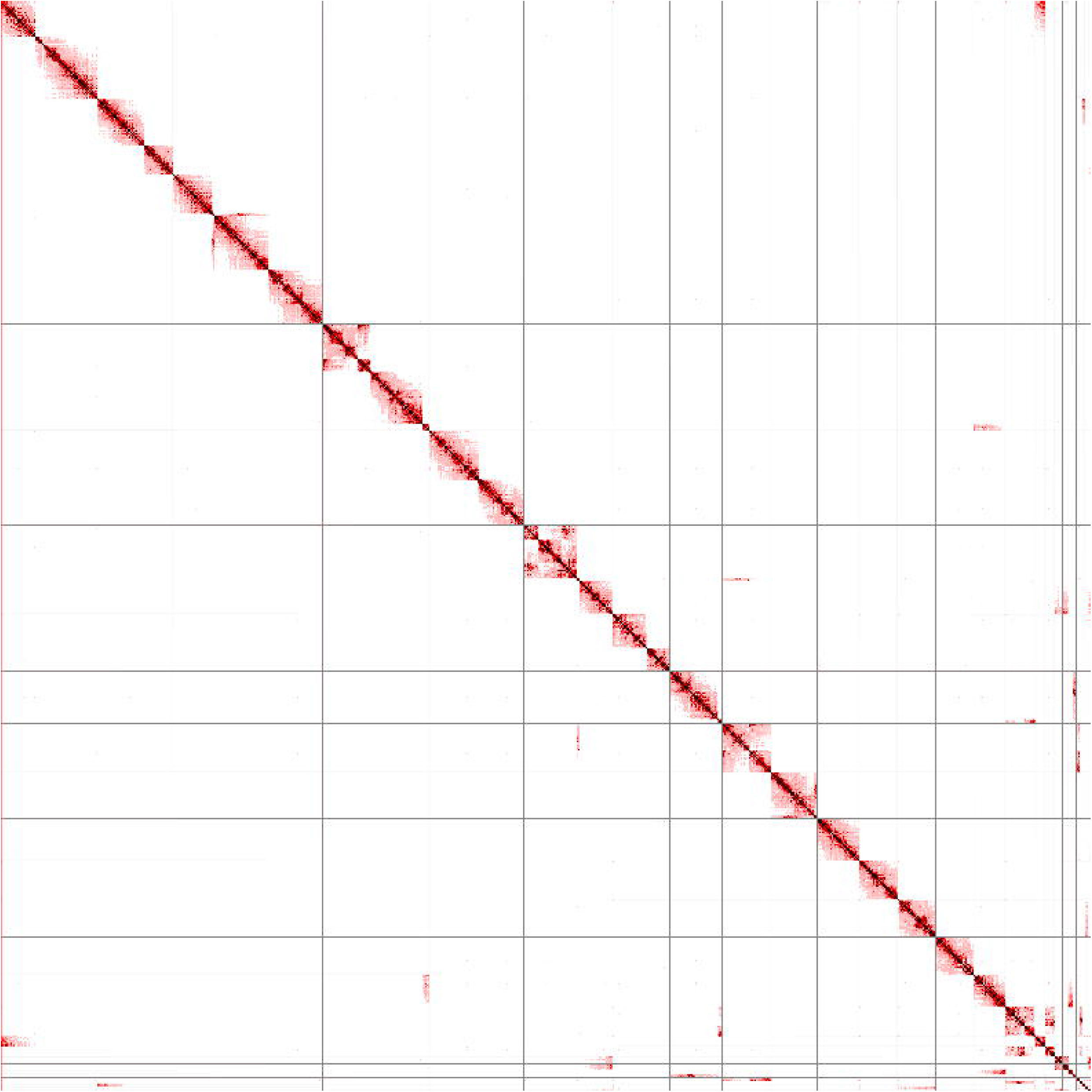
Hi-C Contact Matrix. In the context of scaffolding, Hi-C contact matrices show how correct the scaffolds are based on Hi-C alignment evidence. The longest 26 scaffolds are shown, ordered by descending length from top-left to bottom-right; grey lines show scaffold boundaries. Off-diagonal marks, especially those that are dark and large, are possible evidence of misassembly and/or incorrect scaffolding. Regions with sharp edges similar to where the grey lines appear, but without the grey lines (e.g., three such locations occur in the top-left square), are joins between contigs in that scaffold that lack Hi-C evidence. The lack of Hi-C alignment evidence could suggest that these joins are invalid, but evidence for these joins does exist from RNA-seq alignments. Detection of any spurious joins would, at a minimum, require manual curation. Such curation would enable additional adjustments that would fix minor issues evident from the contact matrix.

The assembly completeness, as assessed with single-copy orthologs, was also evaluated at the contig and scaffold level (Table 4). The results suggest that the modifications made to the primary contig assembly from scaffolding did not significantly impact the complete assembly of single-copy orthologs. The final set of scaffolds had 3,546 complete single-copy orthologs (97.4% of 3,640 from the OrthoDB10 Actinopterygii set). Of these 85.7% (3,120) were present in the assembly only once, and 11.7% (426) were present more than once. Twelve (0.3%) and 82 (2.3%) single-copy orthologs were fragmented in and missing from the assembly, respectively. Approximately 16.7% of the genome was comprised of repetitive elements (Table 5), which is similar to other Carangoid genomes, e.g., *Caranx melampygus* at 16.9% ^11^, *Pseudocaranx georgianus* at 12.8% ^43^, and *Trachinotus ovatus* at 20.3% ^38^.

**Table 4.**
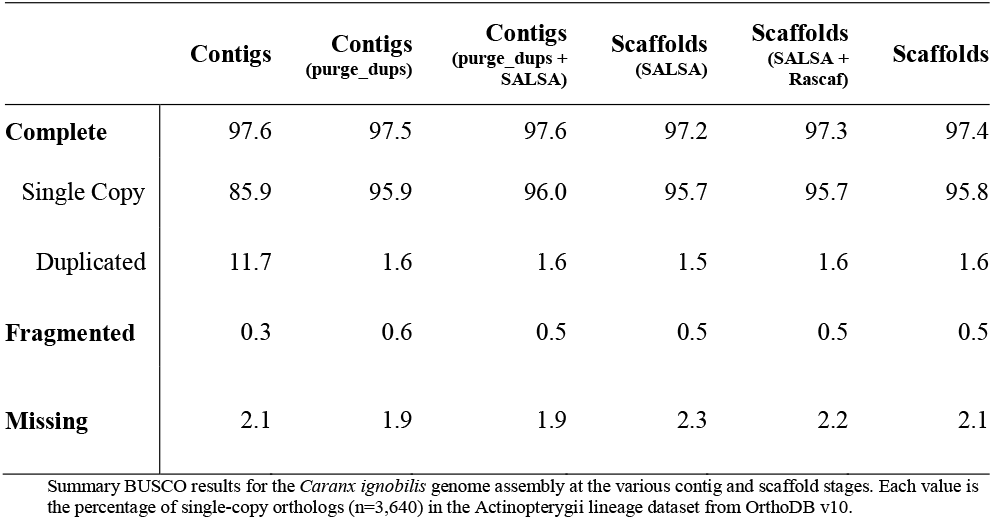
Summary BUSCO Results.

**Table 5.** Summary of Repeats.

|  | **Copies** | **Length**<br>**(Mbp)** | **Percent (%)**<br>**of Sequence** |
| --- | --- | --- | --- |
| **<u>Interspersed Repeats</u>** | **512,100** | **74.5** | **12.3** |
| SINE: | 14,087 | 1.6 | 0.3 |
| Penelope | 5,290 | 1.0 | 0.2 |
| LINE | 62,098 | 12.5 | 2.1 |
| LTR | 15,550 | 3.6 | 0.6 |
| DNA Transposon | 237,928 | 33.1 | 5.5 |
| Unclassified | 177,147 | 23.6 | 3.9 |
| **<u>Tandem Repeats</u>** | **475,796** | **19.2** | **3.2** |
| Satellite | 1,163 | 0.2 | 0.0 |
| SSR | 430,819 | 16.7 | 2.8 |
| Low Complexity | 43,814 | 2.3 | 0.4 |
| **Rolling-circles** | **33,931** | **6.6** | **1.1** |
| **Small RNA** | **7,561** | **1.0** | **0.2** |
| **Total** | **1,029,388** | **100.8** | **16.7** |

Summary of repeat content in the *Caranx ignobilis* genome assembly as reported by RepeatMasker<sup>24</sup> using the Dfam v3.3<sup>25</sup> and RepBase RepeatMasker v20181026<sup>26,27</sup> repeat libraries.

### Comparison of Giant Trevally with Other Carangoid Genomes

We compared the *C. ignobilis* genome to published genomes of other carangoids spanning the carangoid phylogeny, including the live sharksucker (*Echeneis naucrates*) ^32,33^, golden pompano (*Trachinotus ovatus*) ^37,38^, yellowtail (*Seriola quinqueradiata*) ^34,35^, longfin yellowtail (*Seriola rivoliana*) ^36^, greater amberjack (*Seriola dumerili*) ^44,45^, Atlantic horse mackerel (*Trachurus trachurus*) ^39–41^, and closely-related bluefin trevally (*Caranx melampygus*) ^11^. We generated dot plots to visualize genome alignments and look for general synteny between the genomes (Fig. 5). Some structural variation can be seen, but overall, there do not appear to be regions of large variation (e.g., inversions, frameshifts) between *C. ignobilis* and other carangoid species. We similarly compared the same assemblies by visualizing the grouping of single-copy orthologs plotted along the assemblies (Fig. 6). Large groupings of orthologs consistently appear together between genomes, suggesting orthology not just between genes, but also between larger genomic regions. However, at this scale and several genomes compared at once, it is difficult to make more refined inferences on the evolution of specific orthologs within Carangoidei. Additional information could be gleaned if all genomes were assembled at chromosome scale and the sequences were ordered based on similarity.

**Figure 5.**
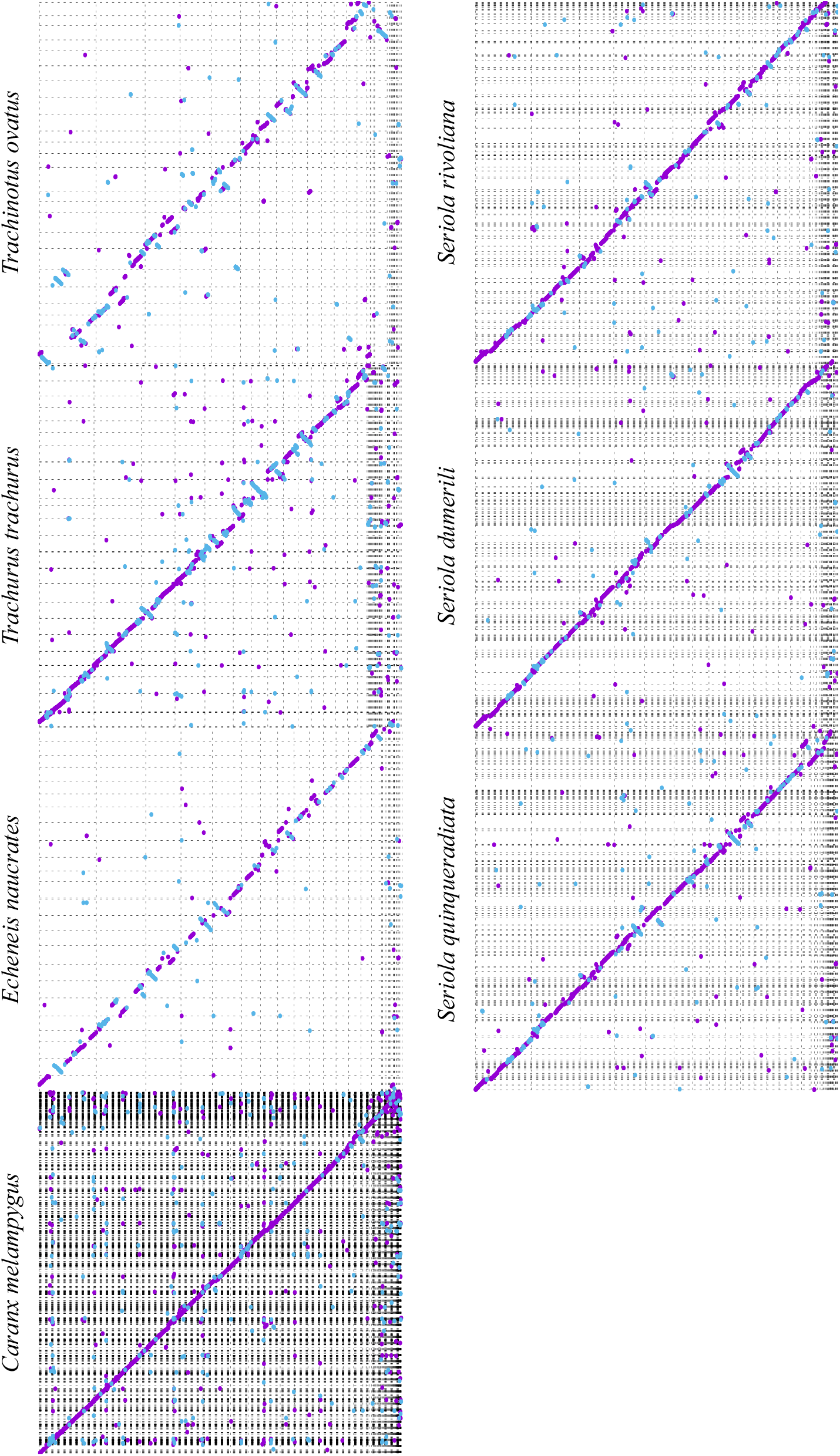
Dot Plot Comparisons with other Carangiformes (Carangoidei) Genomes. Dot plots show the relative continuity of the various segments of two genomes. The purple dots show segments that align in the positive orientation, blue in the negative. The x-axis is the *Caranx ignobilis* genome. The y-axes for each plot are other carangoid genomes. Dots off the diagonal indicate structural variation between the genome assemblies. For assemblies that did not have duplicates purged to reduce the assembly to pseudohaplotypes (*Caranx melampygus* and *Seriola* spp.), the extra dots are presumably the alignment to the secondary copy.

**Figure 6.**
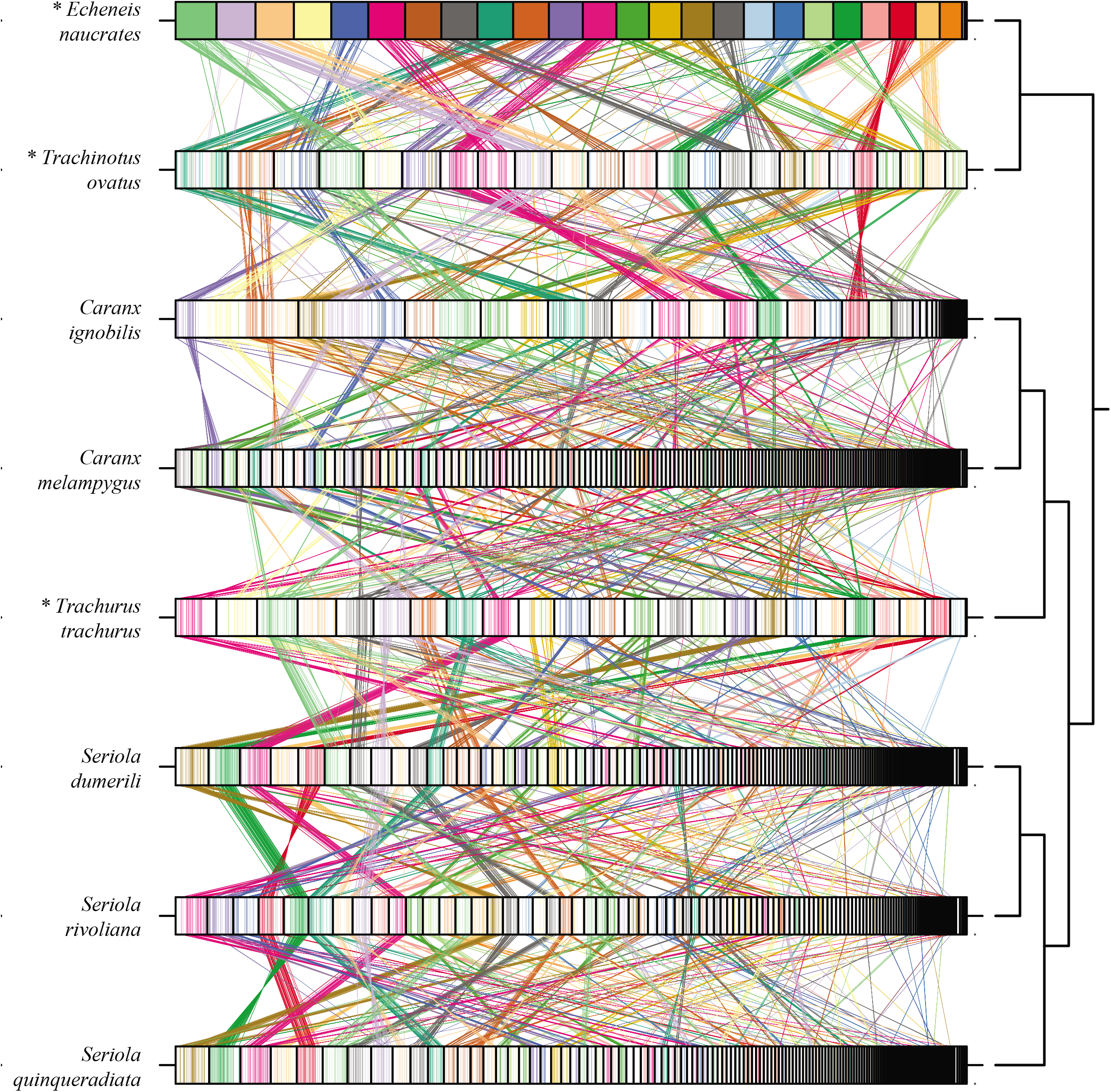
Single-copy Ortholog Comparisons with other Carangiformes (Carangoidei) Fishes. Single-copy orthologs from the Actinopterygii set of OrthoDB v9 were identified with BUSCO v3.0.6 and visualized using ChrOrthLink. “Chromosomes” (usually contigs or scaffolds) are ordered based on length. Note that the sizes of the “chromosomes” are only relative to the other “chromosomes” in the same genome and cannot be compared between genomes. Chromosome-scale assemblies are marked with an asterisk. Colors are assigned based on the *E. naucrates* chromosomes, and individual lines are drawn tracking the placement of individual single-copy orthologs through each genome. Provided that there are no structural rearrangements between different species’ genomes and the genomes are all of reliable quality, large blocks of colored lines should consistently appear together on single chromosomes across the various genomes. Sections of color appearing in blocks on more than one chromosome indicate regions where either chromosome rearrangements occurred or where there were scaffolding errors.

Specific patterns become difficult to inspect at the genome scale when the contigs and scaffolds get small. We observe that the longest scaffolds in the *C. ignobilis* assembly have many single-copy orthologs from more than one chromosome from chromosome-scale assemblies like *E. naucrates*, which calls into question whether some of the *C. ignobilis* scaffolding joins are valid. The joins from Hi-C evidence are trustworthy, but some joins made from RNA-seq data can be spurious under certain conditions — such as when RNA-seq reads split across introns and mapping software mistakenly assigns each end to different genes with similar sequences (e.g., from duplication events or gene families). The true structure of the genome can be further elucidated by karyotype analysis, additional sequencing data (e.g., Ultra-long Nanopore (Oxford, England, UK)), and one-on-one comparisons with high-quality, chromosome-scale assemblies from related species. Ultimately, this genomic dataset is useful for future comparative studies on genome structure and evolution within Carangiformes and marine teleosts more broadly.

## DATA RECORDS

Raw reads have been deposited in the National Center for Biotechnology Information (NCBI) Sequence Read Archive (SRA) ^46–55^ under BioProject PRJNA670456 ^56^, BioSamples SAMN16516519-SAMN16516526 and SAMN16629462 ^57–65^. The genome assembly is associated with the same BioProject under the “container” BioSample SAMN18021194 ^66^ and can be found in GenBank under accession JAFHLA000000000. See Table 6 for a complete list of datasets and their mapping to BioSamples. The contigs, scaffolds resulting from Hi-C evidence, and scaffolds resulting from Hi-C or RNA-seq evidence are also available from the Center for Open Science’s (https://www.cos.io) Open Science Framework ^67^.

**Table 6.** Database Information for Raw Sequences.

| Specimen | Tissue | BioSample Number | Sequencing Type | SRA Accession |
| --- | --- | --- | --- | --- |
| 1 | Blood | SAMN16629462 | Dovetail Omni-C | SRR13036356 |
| 1 | Brain | SAMN16516519 | Illumina RNA-seq | SRR13036363 |
| 1 | Eye | SAMN16516520 | Illumina RNA-seq | SRR13036362 |
| 1 | Fin | SAMN16516521 | Illumina RNA-seq | SRR13036361 |
| 1 | Gill | SAMN16516522 | Illumina RNA-seq | SRR13036360 |
| 1 | Heart | SAMN16516523 | Illumina RNA-seq | SRR13036359 |
| 1 | Heart | SAMN16516523 | PacBio CLR WGA | SRR13036357 |
| 1 | Kidney | SAMN16516524 | Illumina RNA-seq | SRR13036355 |
| 1 | Liver | SAMN16516525 | Illumina RNA-seq | SRR13036354 |
| 1 | Muscle | SAMN16516526 | Illumina RNA-seq | SRR13036353 |

All samples were collected from the same *Caranx ignobilis* specimen in April 2019 off the coast of O'ahu (near Kaneohe, Hawai'i, USA). They are combined under the BioProject PRJNA670456. The genome assembly is deposited in GenBank under accession JAFHLA000000000 with the "container" BioSample SAMN18021194.

## CODE AVAILABILITY

No significant computer programs were generated in this work. Custom scripts referenced in the text are described in the Supplementary Bioinformatics Methods and/or are available on GitHub at https://github.com/pickettbd/caranx-ignobilis_assembly-paper_misc-scripts.

## AUTHOR CONTRIBUTIONS

**JRG:** Funding Acquisition; Writing - Original Draft Preparation; Writing - Review & Editing. **JSKK:** Conceptualization; Funding Acquisition; Investigation; Supervision; Resources; Writing - Review & Editing. **BDP:** Conceptualization; Data Curation; Formal Analysis; Funding Acquisition; Investigation; Methodology; Software; Visualization; Writing - Original Draft Preparation; Writing - Review & Editing. **PGR:** Funding Acquisition; Supervision; Resources; Writing - Review & Editing.

## ACKNOWLEDGEMENTS

We thank the Brigham Young University (BYU; Provo, Utah, USA) DNA Sequencing Center (https://dnasc.byu.edu) and Office of Research Computing (https://rc.byu.edu) for their continued support of our research. We thank the artist, Elaine Heemstra, and the South African Institute for Aquatic Biodiversity (https://www.saiab.ac.za) for the use of the illustration (Fig. 1). The *Trachurus trachurus* genome assembly (GCA_905171665.1) was generated at the Wellcome Sanger Institute as part of the Darwin Tree of Life (DToL; https://www.darwintreeoflife.org) project. We thank Chul Lee of Seoul National University (Seoul, Republic of South Korea) and Ann McCartney and Arang Rhie of the National Institutes of Health – National Human Genome Research Institute (Bethesda, Maryland, USA) for helpful discussion about assembly validation and ChrOrthLink.

