## Supplementary File 1 for "Genome of a Giant (Trevally): *Caranx ignobilis*"

**Supplementary Bioinformatics Methods**

An overview of the methods used in this study was provided in the main manuscript. Where appropriate, additional details, such as the code for custom scripts and the commands used to run software, are provided here.

**Read Error Correction**

The self-corrected reads were generated using Canu v1.8 ^1^ with the following command:

```
canu -correct \
    -s ${SETTINGS_FILE} \
-d ${OUTPUT_DIR_NAME} \
-p ${OUTPUT_PREFIX} \
-pacbio-raw \
${INPUT_PACBIO_READS[@]}
```

The relevant lines of the setting file are included here:

genomeSize=625920000

ovsMethod=sequential

gridEngine=slurm

**Genome Assembly and Scaffolding**

The individual steps of genome assembly and scaffolding will each be described separately. Calculation of assembly summary statistics and the RepeatMasker run will also be described.

*Genome Assembly*

The assembly was created with Canu v1.8 ^1^ using the already corrected reads from the correction process using the following command:

```
canu -trim-assemble \
    -s ${SETTINGS_FILE} \
-d ${OUTPUT_DIR_NAME} \
-p ${OUTPUT_PREFIX} \
-pacbio-corrected \
${INPUT_SELF_CORRECTED_PACBIO_READS_FILE}
```

*Scaffolding and Mis-assembly Detection with Hi-C Data*

Part of the scaffolding process with Hi-C data employed by SALSA is a mis-assembly detection step. The set of contigs created during this process will be pointed out as the scaffolding process is described. The Hi-C data (in this case, Dovetail Genomics Omni-C library using general endonucleases instead of site-specific restriction enzymes) alignments were performed following the Arima Genomics (San Diego, California, USA; https://arimagenomics.com) Mapping Pipeline commit #2e74ea4 (https://github.com/ArimaGenomics/mapping_pipeline), which relied on BWA‑MEM2 v2.1 ^2,3^, Picard v2.19.2 ^4^, and SAMtools v1.9 ^5^. As the pipeline is reasonably well-documented, it will be only summarized here:

1. The assembly (Canu contigs) is indexed using SAMtools faidx.
2. The assembly is indexed with bwa index and the Hi-C reads are mapped to the assembly with bwa mem (I used BWA-MEM2 instead).
3. The alignments are converted from SAM to BAM format with SAMtools view.
4. The 5’ ends are filtered using SAMtools view and the Arima Genomics Perl (https://www.perl.org) script filter_five_end.pl.
5. Paired-end reads are combined into a single file with the Arima Genomics Perl script two_read_bam_combiner.pl and sorted with SAMtools sort. These reads will be treated as single-end hereafter.
6. Read groups are added to the BAM file using Picard AddOrReplaceReadGroups.
7. Merge technical replicates. This step was skipped because no such replicates existed.
8. Duplicates in the BAM file were marked using Picard MarkDuplicates.
9. Merge biological replicates. This step was skipped because no such replicates existed.
10. The final BAM file was indexed with SAMtools index.
11. Stats were reported with the Arima Genomics Perl script get_stats.pl.

Scaffolding was performed on the Canu contigs using the final BAM file from the Arima Genomics Mapping Pipeline with SALSA commit #974589f ^6,7^. First, some pre-processing was required with BEDTools v2.28.0 ^8^ to convert the final BAM file from the mapping pipeline to BED format; this was then sorted. The BEDTools, sorting, and SALSA commands are listed here (note that the ${RESTRICTION_ENZYME_SEQ} was DNASE):

```
bedtools bamtobed \
    -i ${FINAL_ARIMA_BAM_FILE} \
> ${HIC_BED_FILE}

sort -k 4 \
${HIC_BED_FILE} \
> ${SORTED_HIC_BED_FILE}

run_pipeline.py \
-a ${CANU_CONTIGS_FILE} \
-l ${CANU_CONTIGS_FAIDX_FILE} \
-b ${SORTED_HIC_BED_FILE} \
-e ${RESTRICTION_ENZYME_SEQ} \
-s ${GENOME_SIZE} \
-m yes \
-o ${OUTPUT_SALSA_DIR}
```

Note that all newly-created gaps from SALSA will all be assigned a length of 500 nucleotides (i.e., 500 Ns in a row). Assuming these are gaps of unknown size, these will ideally be changed to 100 nucleotides for any submissions to GenBank. If you have multiple sources of evidence for gaps, you will want to keep track of which gaps were supported by each type of evidence. The final command in that set (i.e., run_pipeline.py) iteratively scaffolds with the Hi-C evidence after fixing mis-assemblies. The fixed contigs will be found in a file called assembly.cleaned.fasta and the final iteration of scaffolds will be located in scaffolds_FINAL.fasta. The tiling of contigs (from assembly.cleaned.fasta to create scaffolds_FINAL.fasta) will be in scaffolds_FINAL.agp.

*Scaffolding with RNA-seq Data*

The RNA-seq data were aligned using HiSat v0.1.6-beta ^9^, and the alignments were converted from SAM to BAM format and sorted using SAMtools v1.11 ^5^. First, the assembly (scaffolds from SALSA) was indexed with HiSat. For each tissue (i.e., brain, eye, fin, gill, heart, kidney, liver, and muscle), HiSat aligned reads to the assembly, SAMtools sorted and compressed the output alignments, and Rascaf v1.0.2 commit #690f618 ^10^ computed how scaffolding could be done. The actual scaffolding was done with Rascaf in a single step after all steps had been completed for each tissue. The process is described in the following script:

hisat-build \

${HISAT_IDX_PREFIX} \

${HIC_SCAFFOLDS}

for TISSUE in {brain,eye,fin,gill,heart,kidney,liver,muscle}

do

RNASEQ_READS_LEFT=${TISSUE}_L.fq.gz

RNASEQ_READS_RIGHT=${TISSUE}_R.fq.gz

ALIGNMENT_SAM=${TISSUE}_aln.sam

hisat \

-p ${THREADS} \

--phred33 -q -t \

-x ${HISAT_IDX_PREFIX} \

-1 ${RNASEQ_READS_LEFT} \

-2 ${RNASEQ_READS_RIGHT} \

-S ${ALIGNMENT_SAM}

samtools view \

-buh ${ALIGNMENT_SAM} \

| samtools sort \

-@ ${THREADS} \

-m ${MEMORY}M \

-O BAM \

-o ${ALIGNMENT_BAM}

rascaf \

-breakN 600 \

-b ${ALIGNMENT_BAM} \

-f ${HIC_SCAFFOLDS} \

-o ${TISSUE}.out

done

rascaf-join \

-r brain.out \

-r eye.out \

-r fin.out \

-r gill.out \

-r heart.out \

-r kidney.out \

-r liver.out \

-r muscle.out \

-o ${OUTPUT_FILE_PREFIX}

Note that all newly-created gaps from Rascaf will all be assigned a length of 17 nucleotides (i.e., 17 Ns in a row). Assuming these are gaps of unknown size, these will ideally be changed to 100 nucleotides for any submissions to GenBank. If you have multiple sources of evidence for gaps, you will want to keep track of which gaps were supported by each type of evidence. Also, note that the -breakN option of Rascaf was set to 600 because the gaps from SALSA were 500 bases long. The choice of 600 was arbitrary, it just needed to be longer than 500 (i.e., 501 would have been sufficient). The goal here was to prevent Rascaf from undoing the work SALSA had already done.

Unfortunately, Rascaf does not produce an AGP file like SALSA does. For simplicity in submission to GenBank, such a file is necessary because you would submit the contig-level assembly (contigs made with Canu and fixed with SALSA in this case) and provide an AGP file with scaffold joins and relevant evidence. The information needed to create an AGP file from the Rascaf scaffolds is available in the ancillary output file ending in “.info”. A custom Python script was written to take the contigs file, SALSA AGP file, SALSA scaffolds file, Rascaf scaffolds file, and Rascaf .info file to create two sets of two output files (4 total files). Each set is a fasta and AGP pair where the fasta file is the scaffold level sequence and the AGP file is the description of how to obtain that file from the contig-level file (provided as input). The first set of these files leaves the gaps as they are provided (500 Ns from SALSA and 17 Ns from Rascaf), the second converts them all to 100 Ns. This script is too long to be readable in a document, but the code is available in the file combineHicRna.py on GitHub at https://github.com/pickettbd/caranx-ignobilis_assembly-paper_misc-scripts. During the NCBI submission process, contaminants were identified in the submitted fasta file. These sequences were removed, and appropriate adjustments to the AGP file were also made before resubmission. To create a new scaffold-level fasta file, another custom script was written. It will take an AGP file and input contigs and output scaffolds in fasta format. It is also available in the same GitHub repository in the file agp2fa.py.

*Assembly Statistics*

Assembly continuity statistics, e.g., N50 and auN ^11^, were calculated with caln50 commit #3e1b2be (https://github.com/lh3/calN50) and a custom Python (https://www.python.org) script. caln50 is run using the following simple command:

caln50 \

-s 0.01 \

-L ${GENOME_SIZE} \

${CONTIGS_OR_SCAFFOLDS_FILE} \

> ${STATISTICS_FILE}

The custom Python script is not efficient, but it does calculate Nx, Lx, NGx, and LGx, as well as a few other interesting points about sequences in a fasta file. This script is too long to realistically represent when embedded in the text; it is available on GitHub at https://github.com/pickettbd/basicAsmStatsCalcInPy.

Assembly completeness was assessed using single-copy orthologs with BUSCO v4.0.6 ^12^ and OrthoDB v10 ^13^. The BUSCO config file was the not modified from the default aside from the locations of OrthoDB v10 and the binary executables for BUSCO. It was run based on the following command structure:

busco \

--offline \

--config ${BUSCO_CONFIG_FILE} \

--cpu ${THREADS} \

--in ${CONTIGS_OR_SCAFFOLDS_FASTA} \

--out_path ${OUTPUT_DIR} \

--out ${OUTPUT_FILE_PREFIX} \

--mode genome \

--lineage actinopterygii \

--augustus_species zebrafish

*Repeat Masking*

RepeatMasker v4.1.2-p1 ^14^ was run on the contigs to classify the repeat content of the genome. RepeatMasker was built with dependencies on RMBlast v2.11.0 ^14,15^, TRF v4.09.1 ^16^, hmmer v3.3.2 ^17^, and the h5py (https://www.h5py.org) Python (https://www.python.org) package. Repeat masking was performed against the Dfam v3.3 database ^18^ and the RepeatMasker version of RepBase v20181026 ^19,20^. The command to run RepeatMasker was structured like the following:

RepeatMasker \

-engine "rmblast" \

-species "Carangidae" \

-alignments \

-dir "${OUTPUT_DIR}" \

-html -source -gff -excln \

${CONTIGS_FASTA_FILE}

**Genome Comparisons with Single-copy Orthologs**

Single-copy orthologs were identified from the Actinopterygii set of OrthoDB v9 (same process as for assessing the assembly, but with OrthoDB v9 instead of v10) and BUSCO v3.0.6. These versions of BUSCO and OrthoDB were used, despite being older, because the plotting technique provided by ChrOrthLink depends on the output file structure from BUSCO v3, and BUSCO v4 has changed the format. The commands between BUSCO versions have changed slightly, but they are the same in essence. The command for each genome was based on the following structure:

run_busco.py \

--cpu ${THREADS} \

--in ${ASSEMBLY_FASTA} \

--out_path ${OUTPUT_DIR} \

--out ${OUTPUT_FILE_PREFIX} \

--mode genome \

--lineage_path odb9/actinopterygii \

--species zebrafish

The ChrOrthLink scripts have not yet been prepared for production, so manual editing of the files was necessary to repurpose the code for this analysis. The four scripts (three Python, one R^21^) accept no command-line arguments, so the only way to make it work without adding that functionality is to edit file names and things directly. The simplest way to recreate my analysis or repurpose the ChrOrthLink code for your own analysis in a similar manner would be to clone the repository, edit according to the process described below (substituting your species/filenames/etc. over those described here), and copy your input files into the directory tree. We omitted all sequences that were shorter than 1mb for the plot.

1. Clone the repository. Let’s assume the repo is cloned into a directory called project_dir. Enter the directory and only subdirectory (cd project_dir/VGP_fig5a).
2. Cleanup the stuff you don’t need.

```
rm -rf \
	work/output/* \
	work/BUSCO_genoPlotR_input \
	work/*.csv \
	work/input/BUSCO/*.txt \
	work/input/chr.assign/*.csv \
	work/input/chrsize/*.txt
```

3. Make a note to yourself of some handy abbreviations to use for the genome names. For ours, we used the first letter of the genus and the first 3 letters of the species (e.g., Enau, Cign, Tova, etc.). The rest of these comments will refer to the species name and be meaning this shortened code name as ${SPECIES} (in shell scripts).
4. *Copy* the BUSCO output into work/input/BUSCO. Do not *move* the original output files because these copies will get edited by the scripts; if you made a mistake, it would be annoying to undo the changes when you could have simply re-copied over them. There should be one file per genome included in the analysis (for us, that was eight). The output files from BUSCO are located in the respective BUSCO output directories. The filename is full_table_${SPECIES}.tsv. When copied into work/input/BUSCO, it will need to match the following pattern BUSCO_${SPECIES}.txt.
5. Create the chrsize files. These are formatted as a tab-separated file with the first column being the sequence identifier (from the fasta file, without the >) and the second column being the length of the sequence. The simplest way to obtain this, if you don’t already have it, is to create a fasta index using SAMtools faidx: samtools faidx ${SPECIES}.fa. This will create the index file at ${SPECIES}.fa.fai. The first two columns of this file are what you need. They can be extracted with cut:

```
cut -d \t -f 1-2 \
	path/to/${SPECIES}.fa.fai \
	> work/input/chrsize/chrsize_${SPECIES}.txt
```

6. Create the chr.assign files. These are formatted as comma-separated files with the first column being the sequence identifier (from the fasta file, without the >), the second column being the assigned chromosome number, and the third column being “y” or “n”. If you have curated genomes with assigned chromosome numbers, they can be used. Otherwise, you can make something up. For our plot, we simply assigned chromosome numbers 1-n, where n was the number of sequences in the file. We ordered it based on length of the sequence. We also assigned “y” for the third column for each entry. This can be done with a simple sort and awk command:

```
sort -t \t -n -r -k 2 \
	work/input/chrsize/chrsize_${SPECIES}.txt \
	| awk 'BEGIN{FS="\t"; OFS=",";}{print $1, NR, "y";}'
	> work/input/chr.assign/${SPECIES}.csv
```

7. Edit script #1 in the bin directory (if needed). I changed the location of the work directory, so I had to change the paths, but otherwise this shouldn’t need any fixing. This script will edit the files in work/input/BUSCO and work/input/chrsize based on the files in work/input/chr.assign. Run the script. If a mistake is made when run, you’ll have to re-do steps 4-6 here.
8. Edit script #2 in the bin directory. This script creates *.csv files in work. Change the value of Ref_BUSCO on line 16; we set it to BUSCO_Enau.txt. Run the script.
9. Edit script #3 in the bin directory. This script creates the input for the plot. Change the value of RefID_list on line 19. We set it to ["Enau"]. Change the value of sID_LIST on line 21 to all the species codes. We set it to ["Enau", "Tova", "Cign", "Cmel", "Ttra", "Sdum", "Sriv", "Squi"]. Change the value of target_chr_name on line 23 to "All". I suggest changing the system calls for mkdir around line 570 to include the -p option; this will prevent errors from being unable to create directories that already exist if you re-run these scripts. Run the script.
10. Edit script #4 in the bin directory. Change the value of RefID on line 32. We set it to "Enau". Change the list starting on line 72 to the same names in the same order for sID_LIST as described in step #9. Do the same for the items starting on line 94. Add or remove items for the list starting on line 112 until there are numbers 1-(n-1), with n being the number of species used. In our case, we had 1-7. Run the script. The output should be in work/output. The species names and tree ((("Enau", "Tova"), ((("Cign", "Cmel"), "Ttra"), (("Sdum", "Sriv"), "Squi")));) were manually edited/added in Adobe Illustrator for the final figure.
